## Supplementary material for "ChimpanSEE, ChimpanDO: Grooming and play contagion in chimpanzees": S1 ethogram

### S1 Appendix. Ethogram

*BEHAVIOURS:*

**Grooming**: Picking through the hair of another chimpanzee, searching for and/or removing debris from their body. Involves the use of hands or mouth. May involve any body posture (i.e. sitting, lying down). (Videan et al 2005).

NOTE: does not include genital inspection.
If grooming pauses for <5 secs, ignore the pause, and treat the grooming as continuous.

If grooming stops for >5 secs and <30 secs, record the grooming bout as having ended. The next grooming is a new *grooming bout*.

If grooming stops for >30 secs, or another individual joins the grooming, then any more grooming is a new *grooming episode*.

**Social Play:** Non-aggressive social interaction involving two or more individuals. Never accompanied by pilo-erection. The pattern of behaviour may resemble those used in serious functional contexts (e.g. fighting, mounting, fleeing), but has no obvious immediate benefit to the players (Worch 2012).

- **Low intensity play:**
  Involves low intensity, stationary actions (touching and tickling) (Fröhlich et al 2016), OR any play that is not accompanied by play faces/laughter.
- **Higher intensity play (“Play”):**
  Accompanied by play faces and/or laughter OR involves high-intensity actions (wrestling, chasing, rough and tumble) (Fröhlich et al 2016).

*INITIATION OF BEHAVIOURS:*

An individual initiates a behaviour either if:
(i) they approach another individual and start enacting a behaviour (without their partner performing an initiation gesture)
(ii) they approach another individual and use a particular gesture (see below) which results in the desired behaviour.

The gesture must be an intentional, goal-directed, mechanically ineffective movement, indicated by one or more of the following:

1. Audience checking via eye-gaze
2. Response waiting
3. Sensitivity to recipient’s attentional state
4. Persistence to a goal

**Initiate grooming:** Instigate grooming behaviour through one of the following gestures: (Hobaiter & Byrne; 2011, 2014).

- Big loud scratch: A loud exaggerated scratching movement on the signaller’s own body
- Present grooming: Body is deliberately moved to expose an area to the recipient’s attention, which is immediately followed by grooming of the area
- Bite: recipient’s body is held between the teeth of the signaller

**Initiate play:** Instigate play through on of the following gestures: (Hobaiter & Byrne; 2011, 2014)

(*most common are in italics)*

- Arm shake: Raise arm/hand vertically in the air
- Arm wave: Large repeated back and forth movement of arm raised above the shoulder
- Dangle: To hang from one or both arms from a branch above another individual (audible)
- Drum object palms: Short hard audible contact of alternate palms against an object
- Feet shake: Repeated back and forth movement of feet from the ankles
- *Gallop: An exaggerated running movement where the contact of hands and feet is deliberately audible*
- Hand shake: Repeated back and forth movement of hand from the wrist
- Head nod: Repeated back and forth movement of the head
- *Head stand: Signaller bends forward and places head on the ground*
- Kick: Foot is brought into short hard contact with the recipient’s body in a movement from the hip with a horizontal element
- Knock object: Back of the hand or knuckles are brought into short hard audible contact with an object
- Leg swing: Large back and forth movement of the leg from the hip
- Object in mouth approach: Signaller approaches recipient while carrying an object in the mouth (e.g. a small branch)
- Poke: Firm, brief push of one or more fingers into the recipient’s body
- Pounce: Signaller displaces through the air to land quadrupedally on the body of the recipient
- *Roll over: Signaller rolls onto their back, exposing their stomach, normally accompanied by repeated movements of the arms and/or legs*
- Stomp other: Sole of the foot/feet is lifted vertically and brought into a short had audible contact with the recipient.
