## Supplementary material for "ChimpanSEE, ChimpanDO: Grooming and play contagion in chimpanzees": S2 full results

**S2 Appendix. Full results of GLMMs**

**MODEL 1.1: GROOMING OCCURENCE**

|  |  | Model estimates | | | Chi squared test | |
| --- | --- | --- | --- | --- | --- | --- |
|  | **Term** | **Estimate** | **SE** | **Z** | **chi** | **P (chi)** |
|  | Intercept | -0.691 | 0.462 | -1.496 | *(1)* |  |
| Fixed effects | Age | 0.202 | 0.381 | 0.530 | 0.297 | 0.586 |
|  | Sex | -1.383 | 0.803 | -1.722 | 2.811 | 0.094 |
|  | Social Closeness | 1.550 | 0.846 | 1.833 | 5.904 | 0.015 |
|  | Rank | -0.020 | 0.300 | -0.067 | 0.004 | 0.951 |
| Control effects | No. bouts observed | 0.752 | 0.267 | 2.818 | 9.384 | 0.002 |
|  | No. others | 0.409 | 0.338 | 1.211 | 1.433 | 0.231 |
|  | Time period | -1.319 | 0.692 | -1.906 | 4.255 | 0.039 |
|  | Grooming tendency | 0.660 | 0.319 | 2.068 | 5.100 | 0.024 |

^(1)^ Not indicated due to very limited interpretation

|  |  | Confidence intervals | | Stability tests | |
| --- | --- | --- | --- | --- | --- |
|  | **Term** | **Lower CI** | **Upper CI** | **Min** | **Max** |
|  | Intercept | -2.593 | 0.201 | -0.932 | -0.495 |
| Fixed effects | Age | -0.601 | 1.254 | 0.006 | 0.773 |
|  | Sex | -4.405 | 0.152 | -2.167 | -0.979 |
|  | Social Closeness | 0.344 | 5.535 | 1.217 | 2.295 |
|  | Rank | -0.860 | 0.813 | -0.356 | 0.342 |
| Control effects | No. bouts observed | 0.300 | 1.968 | 0.654 | 1.138 |
|  | No. others | -0.353 | 1.649 | 0.019 | 0.596 |
|  | Time period | -4.255 | 0.068 | -1.978 | -1.086 |
|  | Grooming tendency | 0.084 | 2.025 | 0.521 | 0.915 |

|  | Model estimates | Stability tests | |
| --- | --- | --- | --- |
| Effect | **SD** | **Min** | **Min** |
| Random effect:  Focal ID | 1.112 | 0.863 | 1.519 |
| Random slope:  Focal ID & no. others | 0.782 | 0.350 | 0.956 |
| Random slope:  Focal ID & social closeness | 1.536 | 1.145 | 2.263 |
| Correlation:  Focal ID& no. others | -0.017 | -0.658 | 0.815 |
| Correlation: Focal ID & social closeness | -0.949 | -0.994 | -0.672 |
| Correlation: No. others & social closeness | 0.333 | -0.177 | 0.805 |

**MODEL 1.2: GROOMING LATENCY**

|  |  | Model estimates | | | Chi squared test | |
| --- | --- | --- | --- | --- | --- | --- |
|  | **Term** | **Estimate** | **SE** | **Z** | **chi** | **P (chi)** |
|  | Intercept | -0.683 | 0.193 | -3.529 | *(1)* |  |
| Fixed effects | Age | 0.128 | 0.210 | 0.608 | 0.354 | 0.552 |
|  | Sex | -0.437 | 0.631 | -0.694 | 0.496 | 0.481 |
|  | Social Closeness | -0.270 | 0.151 | -1.790 | 3.116 | 0.078 |
|  | Rank | 0.162 | 0.172 | 0.940 | 0.840 | 0.359 |
| Control effects | No. bouts observed | 0.127 | 0.151 | 0.838 | 0.701 | 0.402 |
|  | No. others | -0.160 | 0.163 | -0.983 | 0.980 | 0.322 |
|  | Time period | -0.920 | 0.523 | -1.760 | 3.290 | 0.070 |
|  | Grooming tendency | -0.095 | 0.183 | -0.518 | 0.268 | 0.605 |

^(1)^ Not indicated due to very limited interpretation

|  |  | Confidence intervals | | Stability tests | |
| --- | --- | --- | --- | --- | --- |
|  | **Term** | **Lower CI** | **Upper CI** | **Min** | **Max** |
|  | Intercept | -1.186 | -0.328 | -0.924 | -0.580 |
| Fixed effects | Age | -0.345 | 0.547 | -0.224 | 0.452 |
|  | Sex | -1.612 | 0.767 | -1.152 | -0.054 |
|  | Social Closeness | -0.614 | 0.000 | -0.338 | -0.196 |
|  | Rank | -0.183 | 0.505 | -0.117 | 0.349 |
| Control effects | No. bouts observed | -0.192 | 0.486 | 0.065 | 0.244 |
|  | No. others | -0.507 | 0.172 | -0.297 | -0.052 |
|  | Time period | -2.072 | 0.085 | -1.145 | -0.540 |
|  | Grooming tendency | -0.519 | 0.275 | -0.173 | 0.026 |

|  | Model estimates | Stability tests | |
| --- | --- | --- | --- |
| Effect | **SD** | **Min** | **Min** |
| Random effect:  Focal ID | 0.096 | 0.000 | 0.528 |
| Random effect:  Event ID | 0.000 | 0.000 | 0.000 |

**MODEL 2.1: PLAY OCCURENCE**

|  |  | Model estimates | | | Chi squared test | |
| --- | --- | --- | --- | --- | --- | --- |
|  | **Term** | **Estimate** | **SE** | **Z** | **chi** | **P (chi)** |
|  | Intercept | -0.831 | 0.354 | -2.344 | *(1)* |  |
| Fixed effects | Age | -1.186 | 0.410 | -2.891 | 11.461 | 0.001 |
|  | Sex | 0.249 | 0.629 | 0.396 | 0.160 | 0.689 |
|  | Social Closeness | 0.454 | 0.303 | 1.500 | 2.649 | 0.104 |
|  | Rank | -0.462 | 0.284 | -1.627 | 2.886 | 0.089 |
| Control effects | No. bouts observed | 0.559 | 0.240 | 2.332 | 5.882 | 0.015 |
|  | No. others | 0.584 | 0.255 | 2.293 | 6.098 | 0.014 |
|  | Time period | -0.293 | 0.513 | -0.570 | 0.328 | 0.567 |
|  | Play tendency | 0.285 | 0.316 | 0.903 | 0.828 | 0.363 |

^(1)^ Not indicated due to very limited interpretation

|  |  | Confidence intervals | | Stability tests | |
| --- | --- | --- | --- | --- | --- |
|  | **Term** | **Lower CI** | **Upper CI** | **Min** | **Max** |
|  | Intercept | -1.836 | -0.235 | -0.977 | -0.636 |
| Fixed effects | Age | -2.387 | -0.513 | -1.638 | -1.021 |
|  | Sex | -1.079 | 1.615 | 0.035 | 0.546 |
|  | Social Closeness | -0.099 | 1.402 | 0.361 | 0.679 |
|  | Rank | -1.139 | 0.072 | -0.625 | -0.340 |
| Control effects | No. bouts observed | 0.119 | 1.210 | 0.235 | 0.673 |
|  | No. others | 0.134 | 1.238 | 0.468 | 0.744 |
|  | Time period | -1.635 | 0.811 | -0.517 | 0.007 |
|  | Play tendency | -0.402 | 1.150 | -0.022 | 0.452 |

|  | Model estimates | Stability tests | |
| --- | --- | --- | --- |
| Effect | **SD** | **Min** | **Min** |
| Random effect:  Focal ID | 0.730 | 0.315 | 0.875 |

**MODEL 2.2: PLAY LATENCY**

|  |  | Model estimates | | | Chi squared test | |
| --- | --- | --- | --- | --- | --- | --- |
|  | **Term** | **Estimate** | **SE** | **Z** | **chi** | **P (chi)** |
|  | Intercept | -1.515 | 0.242 | -6.265 | *(1)* |  |
| Fixed effects | Age | 0.439 | 0.241 | 1.825 | 3.208 | 0.073 |
|  | Sex | -0.276 | 0.354 | -0.780 | 0.615 | 0.433 |
|  | Social Closeness | -0.126 | 0.156 | -0.803 | 0.610 | 0.435 |
|  | Rank | 0.059 | 0.167 | 0.354 | 0.125 | 0.724 |
| Control effects | No. bouts observed | 0.099 | 0.166 | 0.598 | 0.352 | 0.553 |
|  | No. others | 0.186 | 0.183 | 1.020 | 1.008 | 0.315 |
|  | Time period | 0.142 | 0.362 | 0.392 | 0.151 | 0.697 |
|  | Play tendency | 0.077 | 0.202 | 0.379 | 0.142 | 0.706 |

^(1)^ Not indicated due to very limited interpretation

|  |  | Confidence intervals | | Stability tests | |
| --- | --- | --- | --- | --- | --- |
|  | **Term** | **Lower CI** | **Upper CI** | **Min** | **Max** |
|  | Intercept | -2.143 | -1.133 | -1.687 | -1.430 |
| Fixed effects | Age | -0.067 | 1.018 | 0.330 | 0.615 |
|  | Sex | -1.069 | 0.410 | -0.514 | -0.141 |
|  | Social Closeness | -0.493 | 0.228 | -0.227 | -0.058 |
|  | Rank | -0.319 | 0.435 | -0.028 | 0.120 |
| Control effects | No. bouts observed | -0.235 | 0.493 | 0.019 | 0.173 |
|  | No. others | -0.177 | 0.562 | 0.026 | 0.353 |
|  | Time period | -0.588 | 0.976 | -0.097 | 0.342 |
|  | Play tendency | -0.353 | 0.583 | -0.013 | 0.188 |

|  | Model estimates | Stability tests | |
| --- | --- | --- | --- |
| Effect | **SD** | **Min** | **Min** |
| Random effect:  Focal ID | 0.000 | 0.000 | 0.000 |
| Random effect:  Event ID | 0.000 | 0.000 | 0.000 |
